# Regulation of NF-kappa B and cell death by bacterial gingipains

**DOI:** 10.1101/2020.02.25.964585

**Authors:** Sonya Urnowey, Toshihiro Ansai, Vira Bitko, Sailen Barik

## Abstract

The role of non-caspase proteases in the regulation of programmed cell death is relatively under-studied. In this short communication, we report that infection of human gingival fibroblast (HGF) cells *ex vivo* by *Porphyromonas gingivalis* rapidly activated NF-kappa B, an antiapoptotic player; PI3 kinase was also induced, and together, they inhibited apoptosis. Later, pro-apoptotic cellular genes, including caspases, were induced. Apoptosis was evidenced by the degradation of nuclear DNA and activation of caspases. Unexpectedly, a *P. gingivalis* mutant that lacked all three major gingipain proteases was defective in promoting apoptosis, which was consistent with the cleavage of procaspase-3 to the active form by the culture supernatant of wild type *P. gingivalis*, but not that of the triple protease mutant. These results suggest that cellular death in the *P. gingivalis*-infected gum may be orchestrated by gingipain-regulated apoptotic factors.

## 1. Introduction

In periodontitis, the gum (‘gingival’) fibroblast is degraded due to infection and colonization by *Porphyromonas gingivalis* (herein abbreviated as ‘Pg’), a gram-negative anaerobe (Scheres and Crielaard 2013). Several reports have revealed activation of important signaling molecules from Pg-infected cells, which included NF-kappa B (NFκB), IL-1α, IL-8, TNF-α and eotoxin; however, the molecular mechanism of their activation and their roles have been unclear and sometimes conflicting (Amano 2003, Bodet et al. 2005, Brozovic et al. 2006, Diomede et al. 2017, Huang et al. 2001, Kusumoto et al. 2004, Sugano et al. 2004).

*P. gingivalis* famously elaborates a number of proteases, eponymously called ‘gingipains’, (Fitzpatrick et al. 2009, Nakayama 2003, Potempa et al. 2008), which consist of the Arg gingipains, RgpA and RgpB, and the Lys gingipain, Kgp, which cleave Arg-Xaa and Lys-Xaa peptide bonds, and act as major virulence factors in gingivitis (O’Brien-Simpson et al. 2001). Together, they may regulate Pg adhesion and invasion positively or negatively, depending on the cell type (Andrian et al. 2004, Chen et al. 2001, Grenier et al. 2003).

Apoptosis or programmed cell death in the metazoa is controlled by pro- and anti-apoptotic genes, many of which are regulated by intracellular pathogens (Bitko et al. 2007, Guiney 2005, Sinai et al. 2004, Spier et al. 2019). Interestingly, several Pg protease preparations promoted apoptosis when added to cells (Baba et al. 2001, Chen et al. 2001a, Johansson and Kalfas 1998, Morioka et al. 1993, Shah et al. 1992, Sheets et al. 2005, Wang et al. 1999); such ‘extrinsic’ apoptosis could be triggered in part by proteolytic cleavage of cadherins and integrins (Hintermann et al. 2002, Sheets et al. 2005, Tada et al. 2003). Pg infection of human gingival cells of epithelial-type (HGE), on the other hand, triggered an anti-apoptotic pathway, spearheaded by the Bcl-2 protein of mitochondria (Nakhjiri et al. 2001). Yet another study reported rapid mobilization of phosphotidylserine (PS) of the HGE cell membrane following Pg infection, which is a molecular hallmark of an apoptotic response (Yilmaz et al. 2004). Later in infection, the PS was internalized back, coinciding with activated AKT in an anti-apoptotic response. Here, we report the regulation of NF-kappa B and other apoptosis-regulatory factors in Pg-infected primary HGF cells, since these cells are severely affected in clinical gingivitis.

## 2. Experimental methods

Various chemicals and antibodies have been described in the Results section in appropriate places. Essentially all methods have been detailed in previous publications, and thus, only briefly described here. Wild type *P. gingivalis* (Pg) and the triple protease mutant (*rgpA rgpB kgp*) (abbreviately respectively as WT and MT) were grown in anerobic liquid culture media or agar plates (Ansai et al. 1995, Shi et al. 1999). Primary HGF cells were derived from human gingival, obtained from dental clinics with approved IRB, and infected with exponential phase Pg cells washed with phosphate-buffered saline (Lamont et al. 1995, Shi et al. 1999). Bacterial invasion was assessed by antibiotic protection assay (Lamont et al. 1995, Shi et al. 1999). When needed, HGF cells were grown and infected on coverslips, stained with rabbit antibody against Pg Fim A plus TRITC-conjugated secondary antibody, and visualized in a fluorescence microscope (Bitko et al. 2008).

To detect apoptosis, we used multiple complementary assays. Fragmented nuclear DNA was detected by DeadEnd Fluorometric TUNEL assay (Promega, Madison, WI) that coupled fluorescein-12-dUTP to the 3’-OH ends of the DNA, and the labeled DNA was visualized by fluorescence microscopy. The DNA fragmentation assay used the Cell Death Detection ELISA kit (initially from Roche, later Millipore-Sigma). Phosphatidylserine was detected by FITC-conjugated annexin V (Sigma); propidium iodide staining was used to confirm that cells were not necrotic. The fluorescence of the images was also quantified by the NIH ImageJ software (Rabbani et al. 2015).

Luciferase-based assay for functional NFκB quantification has been described (Bitko et al. 1997, Bitko et al. 2004), and where mentioned, appropriate signaling inhibitors were added along with the bacteria.

Caspase-3 activity assay was carried out with the colorimetric substrate, DEVD-pNA, the cleavage of which was detected by increased absorption at 405 nm (Millipore); the reactions were followed in a spectrophotometer cuvette in real time, in order to ensure linearity.

In most graphs, the numbers plotted on the Y-axis are number of folds over uninfected or 0 h or no-enzyme controls, as appropriate. One-way ANOVA and Student’s t-test were used for validation of changes. As routine procedure, all results were from at least three experiments and presented as mean along with error bars for SEM. Protein bands in Western blots (immunoblotting) were developed by the ECL technique (Pierce, Rockford, IL), and have been described in detail before (Bitko and Barik 1998, Bitko et al. 1997).

## 3. Results

### 3.1. Optimization of P. gingivalis infection in cell culture

Past experiments to infect mammalian cell used ∼150-200 bacteria per cell (Lamont et al. 1995), which is clearly a very high multiplicity of infection (m.o.i.). To start with, we queried the role of the high m.o.i. in an effort to optimize the steps of bacterial attachment and invasion, and avoid any non-specific extrinsic effect of the large amount of bacterial material. To quantify the invasion efficiency, washed WT and MT (rgpA rgpB kgp) Pg (Shi et al. 1999) were added to HGF monolayers over a range of m.o.i. (5-200), and the infected cells were counted by indirect immunostaining with Pg antibody (data not shown). The results revealed that only ∼45% HGF cells were infected even at 200 m.o.i., the highest that we tested. Moreover, at all m.o.i, the MT infected only about two-thirds as many cells as did the WT (data not shown). Thus, Pg infects HGF cells rather inefficiently, at least in cell culture, and the gingipains merely improve the efficiency. Based on this optimization, in all subsequent experiments we chose m.o.i. of 80 and 200, respectively, for comparable levels of invasion with the WT and MT bacteria.

When we quantified the progeny Pg released from these infected HGF cells, maximal yield was found at about 72 h post-infection (p.i.) (data not shown) along with lysis of the host cells. For obvious reasons, in analyzing cellular pathways of NF-kB regulation and the apoptotic pathways, we sampled our cells before lysis, i.e., no later than 48 h p.i.

### 3.2. Cell death upon Pg infection

Studies of infection of various gingival cells by *P. gingivalis* have suggested apoptosis by diverse mechanisms that were both dependent and independent of caspases, perhaps owing to differences in cell type and the Pg strain used (Brozovic et al. 2006, Desta and Graves 2007, Sheets et al. 2006). In view of the paramount importance cell death in gingivitis, we argued that the molecular mechanism of Pg-induced apoptosis warrants further studies, which we conducted in our optimized conditions of infection. Kinetics of apoptosis was determined by TUNEL and DNA-fragmentation assays, the results of which agreed with each other and revealed late apoptosis in cells infected with WT Pg, but much less so in those infected with the MT (Fig. 1).

**Fig. 1.**
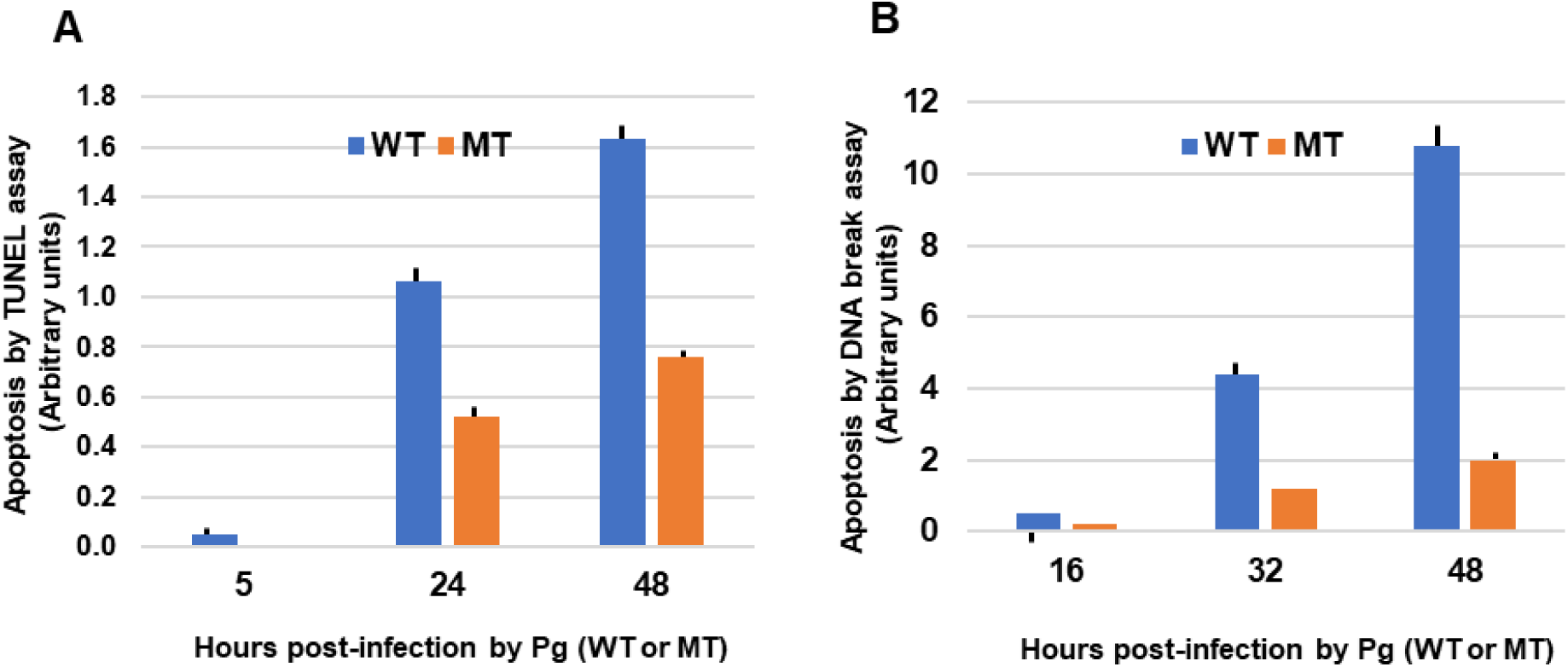
Apoptosis in HGF cells by *P. gingivalis* infection. Cells were infected by wild type (WT) or the triple gingipain mutant (MT), and at the indicated times apoptosis was detected by (A) TUNEL assay and (B) DNA fragmentation ELISA, as described under Experimental Methods.

Apoptosis was found to parallel caspase-3 activation (Fig. 2); necrosis was ruled out (data not shown) by the absence of staining with propidium iodide.

**Fig. 2.**
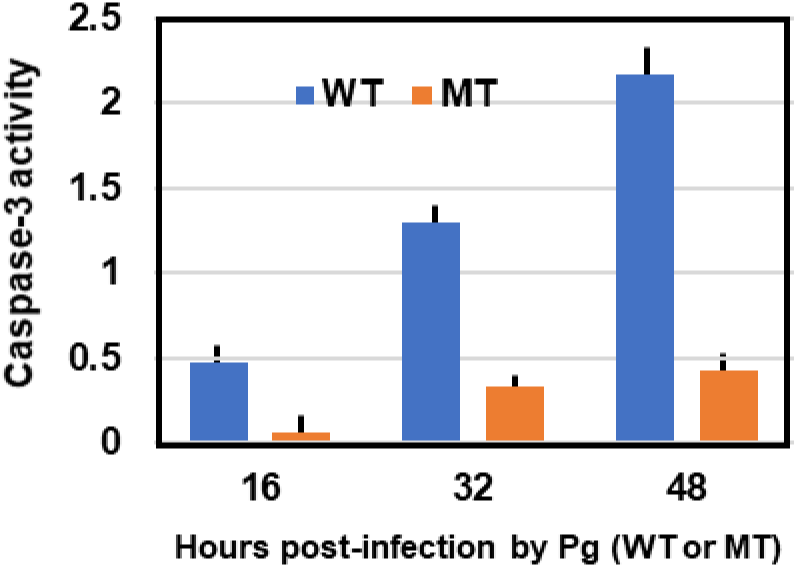
Caspase-3 activation in Pg-infected HGF cells. HGF monolayers were infected with WT or MT Pg, harvested at indicated times, and caspase-3 activity was assayed as described under Experimental Methods. The spectrophotometric A_405_ values are plotted as fold over the uninfected numbers.

We then examined the potential activation of other caspases by immunoblotting with specific antibodies against their cleaved, active fragments (Supplementary material), and the summary results (Fig. 3) not only confirmed the activation of caspase-3, but also those of caspases-6, -7 and -9, but not caspase-8 and -10. Activation of caspases occurred late in infection, starting at ∼24 h, coincident to apoptosis, thus suggesting their role in the apoptosis the Pg-infected cell.

**Fig. 3.**
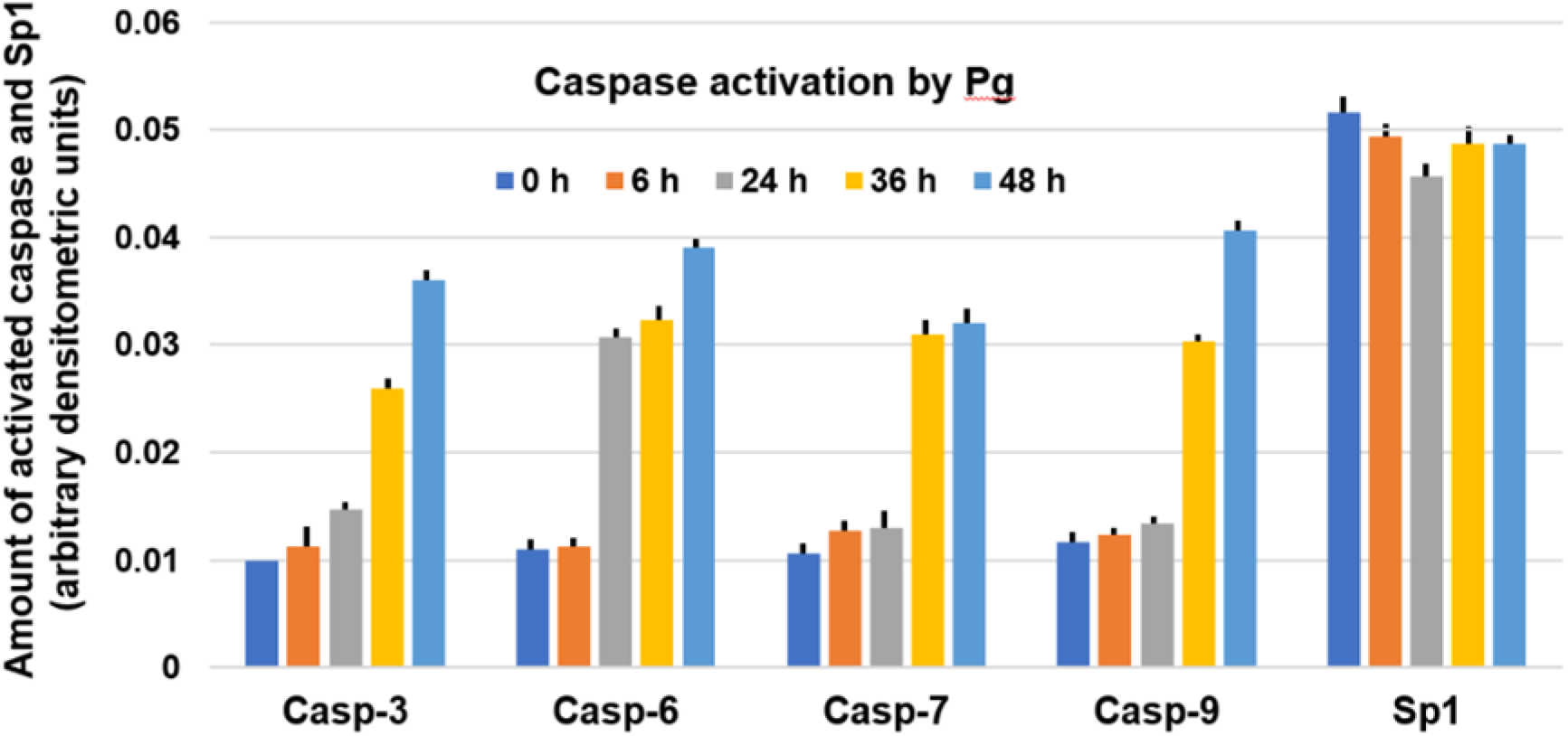
Activation of specific caspases. Pg-infected HGF cells were processed, and active caspase fragments were quantified by immunoblot, followed by densitometry, as described under Experimental Methods.

Lastly, we confirmed intracellular caspase activity in a more direct manner by spectrin-cleavage assay; since caspase-mediated cleavage of the 240 kDa αII-spectrin generates a 120 kDa fragment (Nath et al. 1996, Newcomb et al. 2000). Immunoblot analysis of Pg-infected total HGF extracts indeed showed the fragmentation, but only late in infection, i.e. between 36 h and 48 h (Supplementary material) for both WT and MT Pg.

### 3.3. Procaspase-3 processing activity of Pg proteases

The role of gingipains in invasion is well-documented even though their full impact in host-Pg interaction is not known. Cleavage of cell adhesion molecules have been postulated to contribute to cell death by exogenously added gingipains (Baba et al. 2001, Hintermann et al. 2002, Sheets et al. 2005, Tada et al. 2003, Wang et al. 1999). More recent studies have indicated that non-caspase proteases may also promote apoptosis under certain conditions (Johnson 2000, Neumar et al. 2003, Schrader et al. 2010). Since *P. gingivalis* is laden with proteases, we tested whether they may process procaspase-3 and generate active caspase-3 *in vitro*. Indeed, the supernatant of the WT Pg culture, but not that of the MT, was found to possess potent procaspase-3 processing activity (Fig. 4). Moreover, the cleavage activity was heat-sensitive. In the ‘positive control’ assay, purified caspase-8 did liberate active caspase-3 (Srinivasula et al. 1996). These results suggest that the Arg- and Lys-gingipains of Pg can promote programmed cell death by directly processing intracellular procaspase-3.

**Fig. 4.**
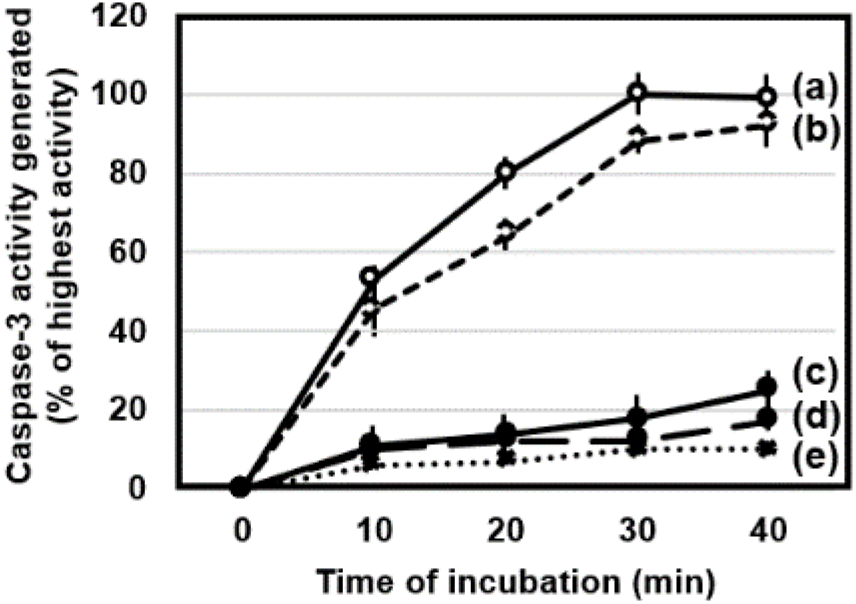
Caspase-3 activation by Pg culture supernatants. Five micrograms of recombinant procsapase-3 (Biomol) substrate was incubated with the following pre-optimized amounts of supernatants / enzymes: (a) WT Pg sup (20 ng); (b) recombinant caspase-8 (4 ng) (Biomol); (c) MT Pg sup (20 ng); (d) WT Pg sup (20 ng), heated at 65 °C, 15 min; (e) sterile growth media. At the indicated times, caspase-3 activity produced in the reaction was assayed as described under Experimental Methods.

### 3.4. NF-kappa B pathway is activated early in Pg-infected cells

With few exceptions (Kucharczak et al. 2003), NF-kappa B (NFκB) functions as an anti-apoptotic factor, and is activated in various infections, whereby it protects the infected cells from premature apoptosis, promoting higher parasite yield (Ansai et al. 1995, Baba et al. 2001, Bitko and Barik 1998; Bitko et al. 1997, Bitko et al. 2007, Chen et al. 2001a, Hintermann et al. 2002, Johansson and Kalfas 1998, Lamont et al. 1995, Morioka et al. 1993, Nakhjiri et al. 2001, Shah et al. 1992, Sheets et al. 2005, Shi et al. 1999, Sinai et al. 2004, Tada et al. 2003, Wang et al. 1999, Yilmaz et al. 2004).

In the classical mechanism of NF-κB activation (Hayden and Ghosh 2004), the cytoplasmic IκB (‘inhibitor of κB’) is phosphorylated by activating signals and dissociates from the p65 (NF-κB) subunit; the latter then translocates to the nucleus, and activates its target genes. Since we noticed a lag in apoptosis in the early stages of Pg infection, we wanted to test the status of NFκB activation in these cells (Fig. 5, next page). This was first observed by NF-κB-dependent reporter luciferase assay (Fig. 5A) as we had previously described for an RNA virus infection (Bitko and Barik 1998, Bitko et al. 2004). In many cells, an upstream phosphoinositol-3 kinase (PI3K) pathway leads to activation of NFκB, and in such cases, the PI3K inhibitor, LY294002, inhibits IκBα phosphorylation (Downward 2004), which we inquired next. First, the PI3K inhibitor (LY294002) (Lin et al. 1995) as well as SN50, the NF-κB inhibitor (data not shown), inhibited luciferase induction. Second, in HGF cells, which are natural hosts of Pg, we confirmed the classical mechanism, i.e. phosphorylation of IκBα (Fig. 5B), followed by localization of p65 in the nuclear fraction (Fig. 5C and Supplementary material), both of which were also inhibited by LY294002 (Fig 5BC). Together, results in both cell types (Fig. 5ABC) documented NF-κB activation relatively early in WT Pg infection, i.e. at ∼15 h, and also revealed that the MT Pg was weaker in this regard. In the PI3K pathway proceeds through a series of cascade phosphorylation and activation, whereby PI3K activates the protein kinase PDK, which in turn activates AKT, a pro-survival kinase. Ultimately, AKT phosphorylates substrates such as forkhead transcription factor FOXO3, IKK, Bad, to name a few. As a result, several anti-apoptotic factors are activated and pro-apoptotic factors (e.g. Bad) are inactivated (Downward 2004, Elmore 2007). We, therefore, determined the phosphorylation status of these signaling molecules in HGF cells during Pg growth, using phosphospecific antibodies. Indeed, several major players of the pathway (AKT, PDK, FOXO3, GSK-3) were found to be phosphorylated early in the HGF cells infected by WT as well as MT Pg (Supplementary material).

**Fig. 5.**
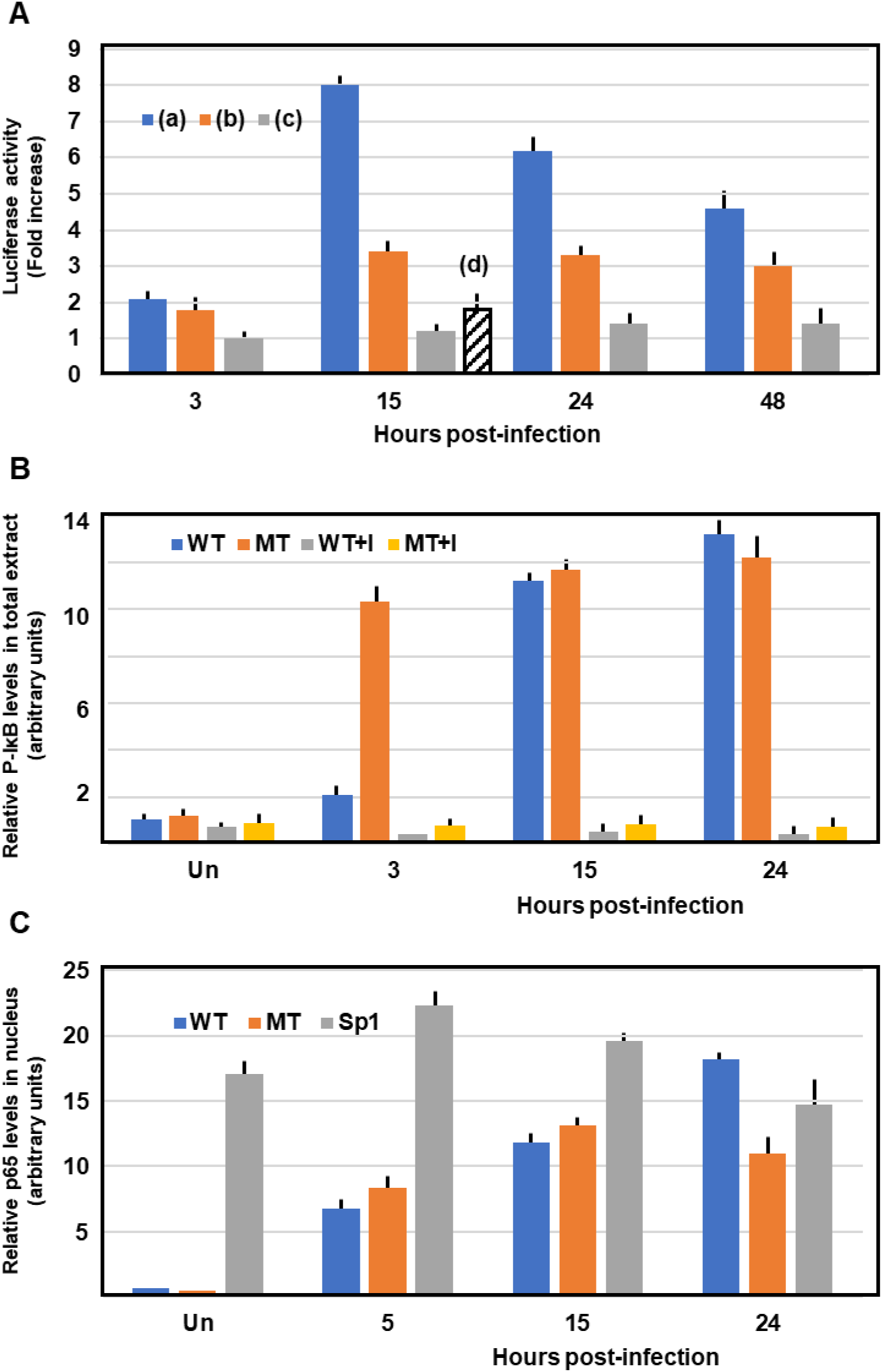
NFκB activation by *P. gingivalis* requires PI3K. (A) NFκB-dependent luciferase activity in Pg-infected engineered HEK cells (Bitko and Barik 1998). Monolayers were infected with wild type *P. gingivalis* in the presence of (a) no drug, (b) 20 μM SN50 (Calbiochem), (d) 20 μM LY294002 (also from Calbiochem; data shown for the 15 h time point only, when the luciferase activity was at its peak, so that inhibition could be easily appreciated), or (c) with the triple gingipain mutant; luciferase assay was then carried out as described in Experimental Methods. (B) Phosphorylated IκBα was detected by immunoblot of the total infected HGF cell extract using a specific antibody (Bitko and Barik 1998). This phosphorylation is essentially undetectable when LY294002 (20 μM) was included in the culture medium (+I). (C) Nuclear import of NFκB p65 subunit. HGF cells were infected with WT and MT *P. gingivalis*, and p65 in 40 μg of nuclear extracts were detected by semi-quantitative by immunoblot (Bitko and Barik 1998, Bitko et al. 2004). Sp1 served as an unchanged control. ‘Un’ indicates uninfected cells.

### 3.5. The apoptotic landscape in Pg-infected HGF cells

Since the results presented above showed activation of an early survival and late apoptotic response following Pg-infection, we then screened a selected cohort of cellular genes that are critical players in the apoptotic commitment and decision (Elmore 2007, Riedl and Shi 2004; Siddiqui et al. 2015). Transcriptional induction of pro- and anti-apoptotic genes were quantified by qRT-PCR using commercially made primers, and representative results (Fig. 6) revealed a trend of early induction of anti-apoptotic (i.e. pro-survival) genes and late induction of pro-apoptotic ones (Bock and Tait 2020, González-Arzola et al. 2019, Ow et al. 2008).

**Fig. 6.**
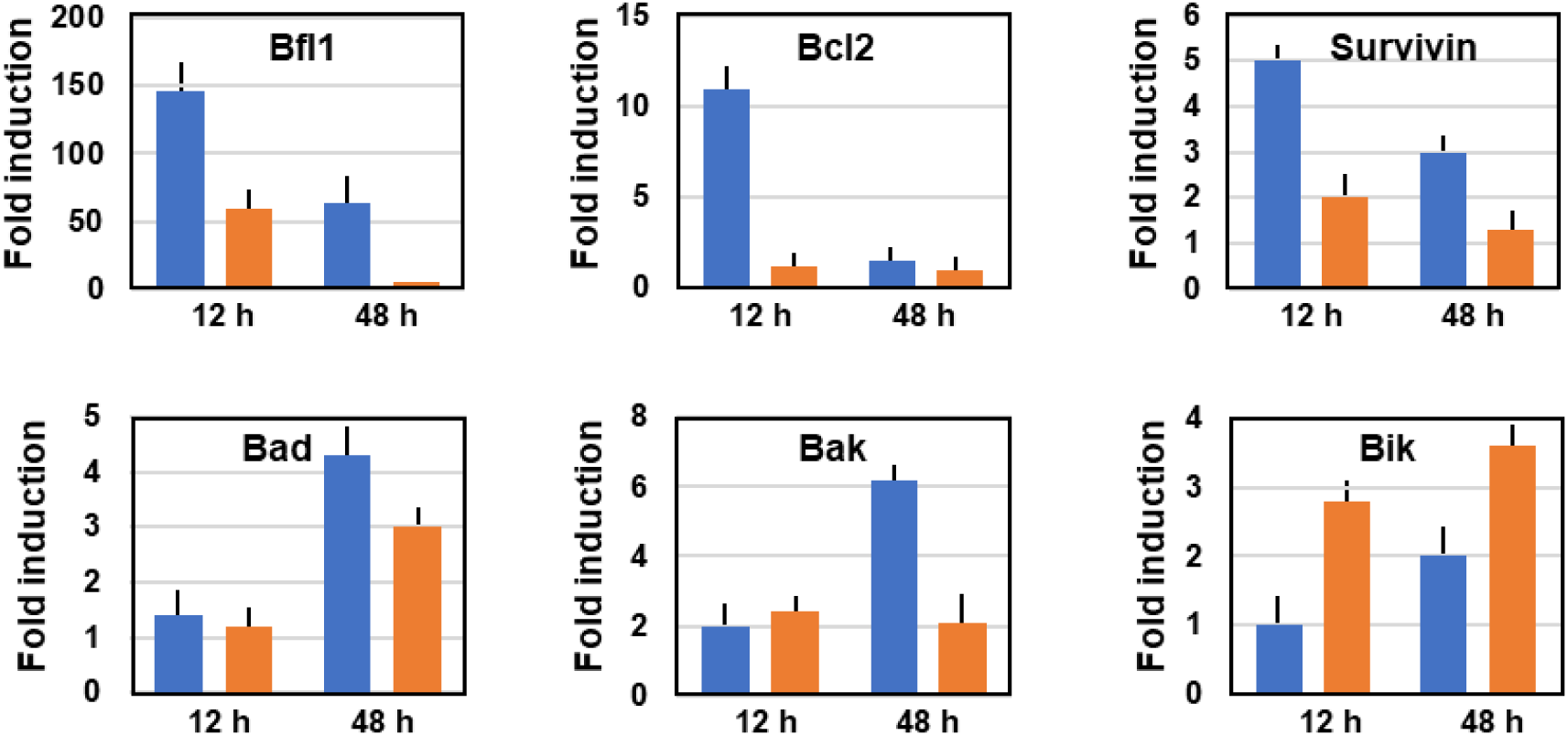
The kinetics of apoptotic gene regulation in Pg infection. Quantitative real-time PCR (qRT-PCR) was performed as described (Bitko et al. 2004). Fold induction over the uninfected cell value is plotted for 12 h and 48 h post-infection by WT (Blue bars) and MT (Orange bars) Pg. Three cardinal anti-apoptotic (Bfl2, Bcl2, survivin) and three pro-apoptotic (Bad, Bak, Bik) genes are shown as class representatives.

LY294002 and SN50, which are inhibitors of PI3K and NFκB, respectively (Lin et al. 1995), suppressed the induction of several anti-apoptotic genes (data not shown to save space). These results further correlate the anti-apoptotic gene expression profile with the activation of the survival pathway at early times of Pg infection.

## 4. Discussion

Results presented here are in accord with previous reports suggestive of an inefficient interaction of Pg with host fibronectin / integrin network (Njoroge et al. 1997, Umemoto and Hamada 2003, Yilmaz et al. 2002) and improvement of such interactions by Arg-gingipains (Kontani et al. 1997), which implied that the cellular receptors of Pg are conformationally exposed by the Arg-gingipains. The high amounts of Pg required to produce a decent percentage of infected cells is likely because such interactions are naturally inefficient and require a higher bacterial load. The lower efficiency of invasion by the gingipain mutant supports this mechanism. The significance of the inefficient invasion process is currently a matter of conjecture.

Substantial amount of evidence presented here, which included kinetics of activation of the relevant factors, has clearly revealed a sequential nature of the apoptotic program in the Pg-infected cell, whereby the survival response is triggered first, apoptotic response later. The advantage to Pg is easier to explain in this scenario, since as mentioned before, prolonging the cellular integrity should allow a longer growth cycle of the intracellular bacteria. It may also shield the bacteria from the immunity of the host. However, it is unclear, whether the host derives any evolutionary benefit in this order of interaction. The late-stage apoptosis and physical lysis of the host is necessary for bacterial egress, and the role of gingipains in this process is easier to postulate because of their ability to activate caspase-3. It is much harder to conceive a role of the gingipains in the survival phase, since proteases are degradative in nature. Perhaps the gingipains promote survival by degrading a negative regulator of the survival pathway, such as proapoptotic factor; alternatively, if they may function in a nonenzymatic role, perhaps by association with a death-related signaling protein. In any event, this remains a fascinating area to study in the future.

Pioneering early studies (Eichinger et al. 1999) revealed a ‘caspase-like’ fold in the Arg-gingipain, but its physiological relevance remained unresolved. Subsequent sequence homology studies suggested a ‘caspase-hemoglobinase’ fold in several members of the protease superfamily, which included human legumain and bacterial clostripain (clostripeptisae B, from *Clostridium histolyticum*) and Pg gingipains (Aravind and Koonin 2002, Barrett and Rawlings 2001, Chen et al. 1998, Mikolajczyk et al. 2003). It will be interesting to determine if the cleavage in Pg-activated caspase-3 occurs near the natural processing site (Cohen 1997).

## Acknowledgments

The *P. gingivalis* strains were kindly provided by Professor Koji Nakayama, Division of Microbiology and Oral Infection, Nagasaki University Graduate School of Biomedical Sciences, Sakamoto 1-7-1, Nagasaki 852-8588, Japan. Dr. Tadamichi Takehara in the Department of Preventive Dentistry, Kyushu Dental College, provided initial support to TA for this project. This research was funded in part by National Institute of Health (grants AI045803, EY013826), USA.

