## Supplemental file for "Regulation of NF-kappa B and cell death by bacterial gingipains"

**<u>Supplementary Material</u>**

**Supplementary data 1.** (A) Activated (cleaved) caspases and control Sp1 protein in the cell extract were detected by immunoblot using primary antibodies from Santa Cruz Biotechnology. Note that caspases-8 and -10 were different from the other caspases for not being activated. (B) Full-length of α-spectrin (240 kDa) and its caspase-generated fragment (120 kDa) in the same extracts were detected by antibody from Chemicon International.

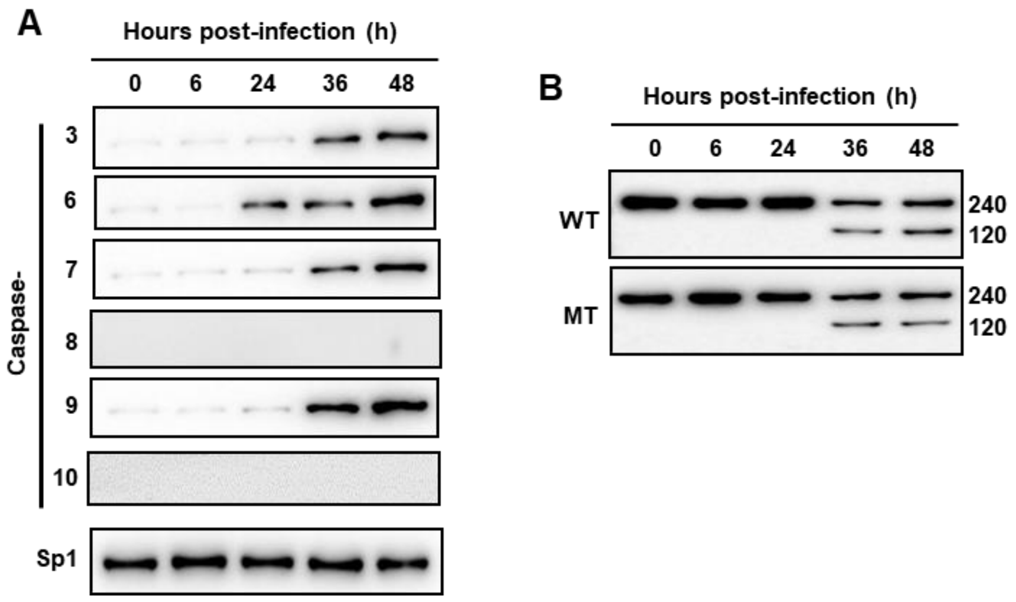


**Supplementary data 2.** Immunoblot of (A) total cell phospho-IκBα and (B) nuclear p65 (NFκB) at different times after Pg infection of HGF cells. ‘Un’ indicates uninfected cells, ‘+I’ means in the presence of LY294002.

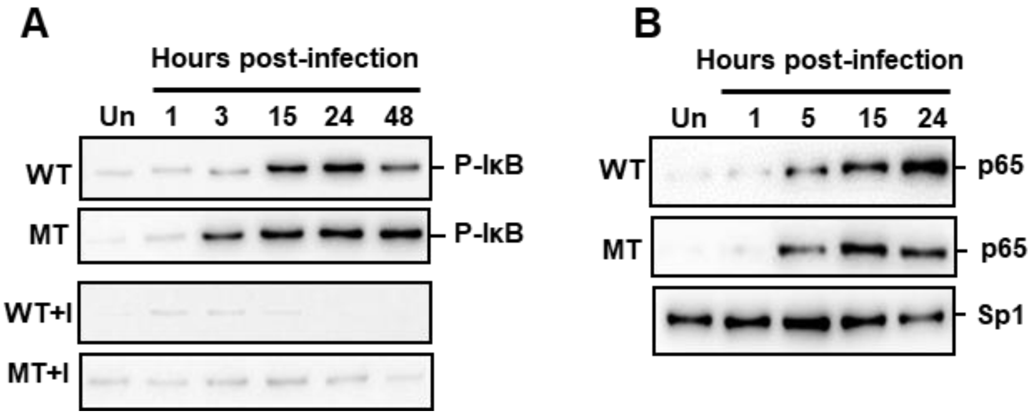

**Supplementary data 3.** Signaling through the PI3K/AKT pathway in Pg-infected cells. HGF monolayer was infected with WT or MT Pg bacteria as before, and the infected cells were grown further in the presence (+I) or absence of 20 μM LY294002 (PI3K inhibitor). Cell extracts (50 μg), made in the presence of phosphatase inhibitor mix (Calbiochem), were probed in immunoblot for total AKT or specific phosphorylated proteins of the AKT pathway using antibodies from New England Biolabs. The two species of phospho-FOXO3 can be seen. Note the early (6-12 h) activation of phosphorylation of all proteins and its inhibition by LY294002.

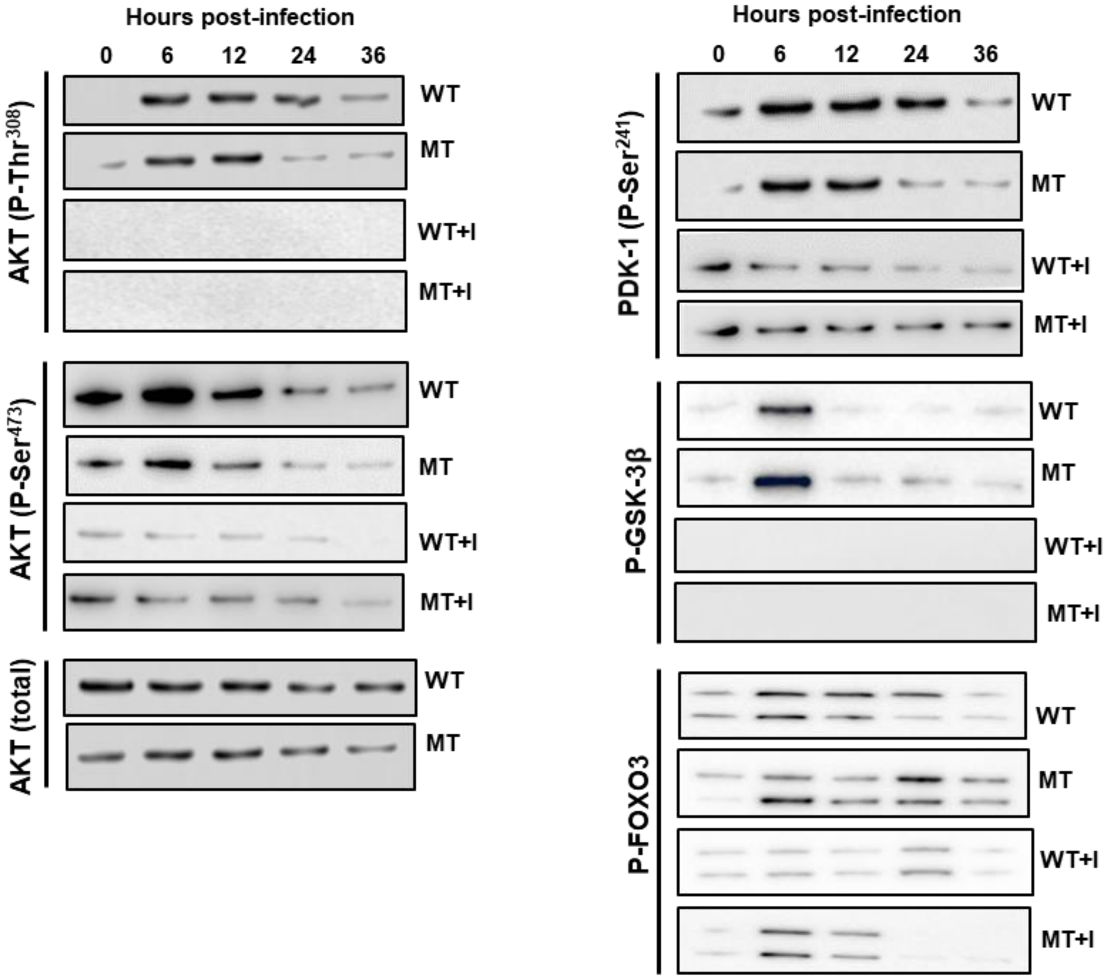
